# Discovery of tandem and interspersed segmental duplications using high throughput sequencing

**DOI:** 10.1101/393694

**Authors:** Arda Soylev, Thong Le, Hajar Amini, Can Alkan, Fereydoun Hormozdiari

## Abstract

**Motivation:** Several algorithms have been developed that use high throughput sequencing technology to characterize structural variations. Most of the existing approaches focus on detecting relatively simple types of SVs such as insertions, deletions, and short inversions. In fact, complex SVs are of crucial importance and several have been associated with genomic disorders. To better understand the contribution of complex SVs to human disease, we need new algorithms to accurately discover and genotype such variants. Additionally, due to similar sequencing signatures, inverted duplications or gene conversion events that include inverted segmental duplications are often characterized as simple inversions; and duplications and gene conversions in direct orientation may be called as simple deletions. Therefore, there is still a need for accurate algorithms to fully characterize complex SVs and thus improve calling accuracy of more simple variants.

**Results:** We developed novel algorithms to accurately characterize tandem, direct and inverted interspersed segmental duplications using short read whole genome sequencing data sets. We integrated these methods to our TARDIS tool, which is now capable of detecting various types of SVs using multiple sequence signatures such as read pair, read depth and split read. We evaluated the prediction performance of our algorithms through several experiments using both simulated and real data sets. In the simulation experiments, using a 30× coverage TARDIS achieved 96% sensitivity with only 4% false discovery rate. For experiments that involve real data, we used two haploid genomes (CHM1 and CHM13) and one human genome (NA12878) from the Illumina Platinum Genomes set. Comparison of our results with orthogonal PacBio call sets from the same genomes revealed higher accuracy for TARDIS than state of the art methods. Furthermore, we showed a surprisingly low false discovery rate of our approach for discovery of tandem, direct and inverted interspersed segmental duplications prediction on CHM1 (less than 5% for the top 50 predictions).

**Availability:** TARDIS source code is available at https://github.com/BilkentCompGen/tardis, and a corresponding Docker image is available at https://hub.docker.com/r/alkanlab/tardis/

## 1 Introduction

Genomic differences between individuals of the same species, or among different species, range from single nucleotide variation (SNVs) [22] to small insertion/deletions (indels) [26] up to 50 bp, structural variation (SVs) [2] that affect >50 bp, and larger chromosomal aberrations [28]. Among these types of variants, SNVs were extensively and systematically studied since the introduction of microarrays, which can also be used to genotype short indels [22]. SVs, especially copy number variations (CNVs), were first identified using BAC arrays [33, 31], and then oligonucleotide array comparative genomic hybridization [34, 9] and SNV microarrays by analyzing allele frequencies[23, 10]. Chromosomal aberrations such as trisomy, or large translocations (e.g., Philadelphia chromosome [32]) can be tested using fluorescent in-situ hybridization [28].

Fine scale SV discovery was made possible using fosmid-end sequencing [44], and later indels were identified at breakpoint level using whole genome shotgun (WGS) sequencing data [26]. However, both approaches used the Sanger sequencing technology, which is prohibitively expensive to scale to analyze thousands of genomes. High throughput sequencing arose as a cost effective alternative [35] to characterize SVs first using the Roche/454 platform [18], and then Illumina [3, 12, 45, 25, 20, 36, 1, 45].

The 1000 Genomes Project, launched in 2008, used the HTS platforms to catalog SNVs, indels, and SVs in the genomes of 2,504 human individuals [41]. Many algorithms were developed that use one of four basic sequence signatures to discover SVs, namely read depth, read pair, split reads, and assembly [24, 2], however, most of these tools focus on characterizing only a few types of SVs. More modern SV callers such as DELLY [30], LUMPY [19], SV-Bay [17], TIDDIT [11], and TARDIS [37] integrate multiple sequencing signatures to identify a broader range of SVs such as deletions, novel insertions, inversions, and mobile element insertions. However, there is still a need for accurate algorithms to characterize several forms of complex SVs, such as tandem or interspersed segmental duplications (SDs) [8, 7]. Note that read depth based methods can identify the *existence* of SDs [3, 39], but cannot detect the location of the new copies of the duplications. Only SV-Bay [17] and TIDDIT [11] are capable of reporting duplication insertion location using read pair information.

Here we describe novel algorithms to accurately characterize both tandem and interspersed SDs using short read HTS data. Our algorithms make use of multiple sequence signatures to find approximate locations for the duplication insertion breakpoints. We integrated our methods into the TARDIS tool [37] therefore extending its capability to simultaneously detect various types of SVs. We test the new version of TARDIS using both simulated and real data sets. We show that TARDIS achieves 96% sensitivity with only 4% false discovery rate (FDR) in simulation experiments. We also used real WGS data sets generated from two haploid genomes (i.e., CHM1 [15] and CHM13 [38]). Comparison of our predictions with *de novo* assemblies generated using long reads from the same DNA resources [38] revealed ¡5% false discovery rate for the duplications with high score.

The algorithms we describe in this manuscript are among the *first* methods to discover the insertion locations of segmental duplications using high throughput sequencing data. Coupled with the previously documented capability of TARDIS to identify deletions, novel and mobile element insertions, and inversions, we are one more step closer towards a comprehensive characterization of SVs in high throughput sequenced genomes.

## 2 Methods

### 2.1 Motivation

The 1000 Genomes Project provides a catalog of SVs in the genomes of 2,504 individuals from many populations [40]. The project primarily focused on characterizing deletions, insertions, and mobile element transpositions, however, it also generated a set of inversion calls. A careful analysis shows that a substantial fraction of the predicted inversions are in fact complex rearrangements that include duplications, inverted duplications, and deletions within an inverted segment (Figure 1). This is because the read pair signatures that signal such complex SVs are exactly the same as shown in Fig. 2. Therefore, any algorithm based on read pair (and/or split read) signature may incorrectly classify these complex events as simple inversions, unless it tries to characterize all such events simultaneously, with additional probabilistic models to differentiate events that show themselves with the same signature.

**Figure 1:**
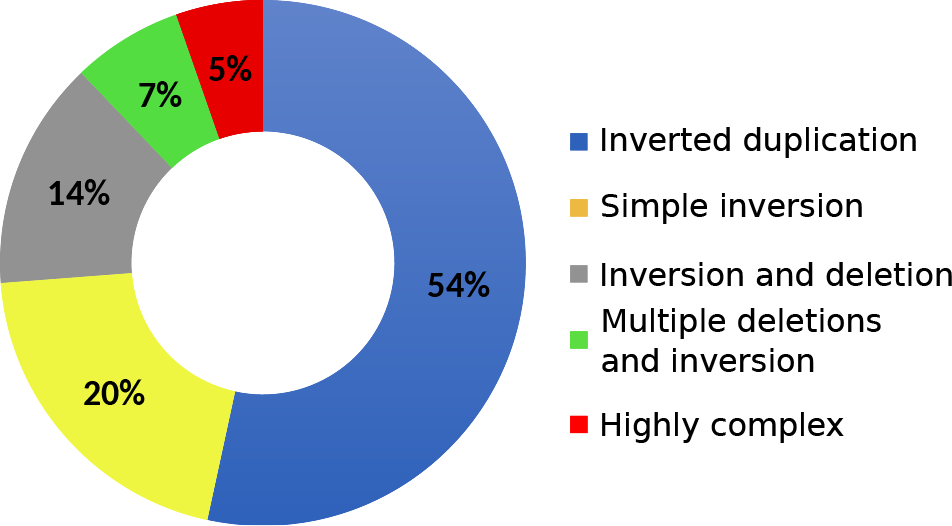
Relative abundance of complex SVs among the inversion calls reported in the 1000 Genomes Project [40]. 54% of predicted inversions are in fact inverted duplications and only 20% are correctly predicted as simple inversions.

**Figure 2:**
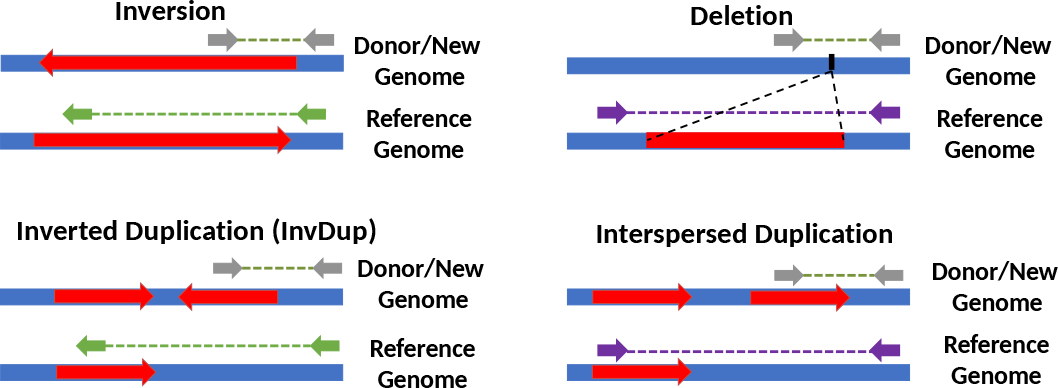
Read pair sequence signatures of inversions, deletions and segmental duplications. The grey arrows show read pairs that span a structural variant breakpoint, and green (left panel) and purple (right panel) arrows show the corresponding map location and orientation of these reads on the reference genome. Note that the read pair signatures for inversions and inverted duplications are exactly the same. Similarly, deletions and direct duplications show the same read pair signature. Therefore read pair based algorithms may incorrectly identify inverted segmental duplications as simple inversions. This problem also exists for incorrectly predicting simple deletions while the true underlying variant is a duplication in direct orientation.

### 2.2 Read pair and split read clustering

TARDIS uses a combination of read pair, read depth and split read sequencing signatures to discover SVs [37]. TARDIS formulation is based on algorithms we developed earlier using maximum parsimony [12, 14] objective function. The proposed approach has two main steps: First clustering read pairs and split reads that signal each specific type of SV, and second apply a strategy to select a subset of clusters as predicted SV. In this paper we extend TARDIS to characterize a complex set of SVs, which are incorrectly categorized by state of the art methods for SV discovery. Specifically the methods we present here will **advance our capability in discovery of duplication based SVs**. Furthermore, our new methods are capable of separating inversions from more complex events of inverted duplications and are also able to predict the insertion locations of the new copies of segmental duplications. We would argue that considering these more complex types of SV is crucial in improving the accuracy of predicting other types of SVs. We therefore modified TARDIS to calculate a likelihood score for each SV provided the observed read pair, read depth and split read signatures. Figure 3 summarizes the read pair signatures that TARDIS uses to find tandem in direct orientation and interspersed duplications in both direct and inverted orientation. Although not shown on the figure for simplicity, similar rules are required for split reads that signal the same types of SVs (Supplementary Figure 1).

**Figure 3:**
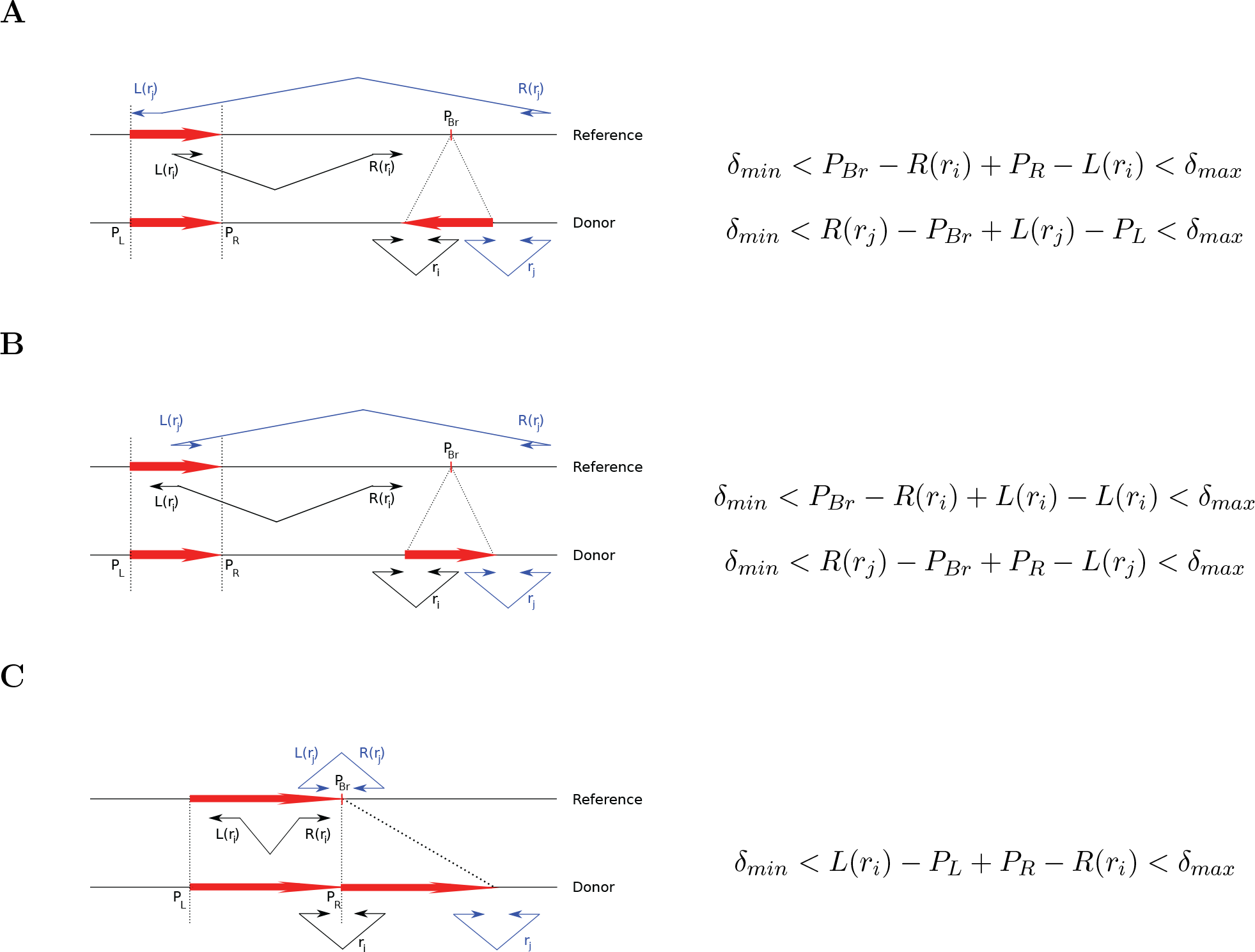
Read pair sequence signatures used in TARDIS to characterize A) interspersed duplications in inverted orientation, B) interspersed duplications in direct orientation, and C) tandem duplications. *P_Br_* denotes the breakpoint location of each variant, and *P_L_* and *P_R_* are the left and right (i.e., proximal and distal) coordinates of the duplicated segment. For each type of structural variation, we show two read pairs from the donor genome (*r_i_, r_j_*). The read pairs are colored black and blue to facilitate easier tracking by the reader. The alignments for read pair *r_i_* are shown on the reference as *L*(*r_i_*) and *R*(*r_i_*), which denote the left (i.e., proximal) and right (i.e., distal) mapping locations of the end reads. Finally, *δ_min_* and *δ_max_* are the minimum and maximum fragment lengths as inferred from the fragment size distribution in the aligned data.

#### 2.2.1 Maximal valid clusters

Our approach for discovery of SVs is based on first produced maximal valid clusters for every type of SVs. We have previously described algorithms to calculate maximal valid clusters for deletions, inversions, and mobile element insertions [12, 13, 14, 37]. A valid cluster is defined as a set of discordant paired-end read alignments that support the same structural variants. In another words, a valid cluster indicates the set of discordant paired-end read mappings that explain the same potential structural variant. More formally, a valid cluster is a set of alignments of discordant read pairs and/or split reads (denoted as *rp_i_*) that support the same particular SV event shown as

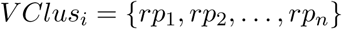

A maximal valid cluster is a valid cluster which no additional discordant paired-end reads can be added to it such that it still remains a valid cluster. Note that, we and others have previously developed methods to efficiently generate all maximal clusters for inversions, deletions, and insertions. In this section we provide new methods to find maximum valid clusters for tandem and interspersed (both direct and inverted) duplications.

There are a set of rules that each *rp_i_* should satisfy in order to support the cluster, *V Clus_i_*, based on the type of SV.

##### Inverted duplications

We assume the fragment sizes for read pairs are in the range [*δ_min_, δ_max_*], and we denote the insertion breakpoint of the duplication as *P_Br_* and the locus of the duplicated sequence is [*P_L_, P_R_*] (Figure 3A). We scan the genome from beginning to end, and we consider each position as a potential duplication insertion breakpoint *P_Br_*. We consider all sets of read pairs where both mates map to the same strand (i.e., +/+ and −/−) within interval [*P_Br_ − δ_max_, P_Br_*] and [*P_Br_, P_Br_* + *δ_max_*] respectively as clusters that potentially signal an inverted duplication.

##### Interspersed direct duplications

We create the valid clusters in a way similar to the inverted duplications, with the exception of the required read mapping properties. For direct duplications we require each mate of a read pair to map to opposing strands (i.e., +/− and −/+).

##### Tandem duplications

We also create the clusters for tandem duplications as shown in Figure 3. In the case of tandem duplications, discordant read pairs and split reads map in opposing strands, where the read mapping to the upstream location will map to the reverse strand, and the read mapping to downstream will map to the forward strand (i.e., −/+).

Similar to the valid cluster formulation, a maximal valid cluster is a valid cluster that encompasses all the valid read pairs and split reads for the particular SV event (i.e., no valid superset exists). This can be computed in polynomial time as follows:

1. We initially create maximal sets *S* = {*S*_1_*, S*_2_*, …, S_k_*} that harbors the read pair/split read alignments *S_i_* = {*rp*_1_*, rp*_2_*, …, rp_k_*}.
2. For interspersed duplications, we use an additional step to bring mappings in both forward-reverse and reverse-forward (forward-forward and reverse-reverse for inverted duplications) orientations together inside the same set.
3. For each maximal overlapping set *S_i_* found in step 1, we create all the overlapping maximal subsets *s_i_*. (This step is necessary only for detecting inversions and interspersed duplications)
4. Among all the sets *s_i_* found in Step 3, remove any set that is a proper subset of another chosen set.

### 2.3 Probabilistic Model

As we describe above different types of SVs may generate similar discordant read pair signatures (Figure 2). We therefore developed a probabilistic model that makes use of the read depth signature to assign a likelihood score to each potential SV. Our new probabilistic model has the ability to distinguish different types of SVs with the same read pair signature.

#### 2.3.1 Likelihood model

Assume the set of maximum valid clusters *SV* = {*S*_1_*, S*_2_*, …, S_n_*} is observed in the sequenced sample. TARDIS keeps track the following information for each maximum valid cluster *S_i_* for 1 *≤ i ≤ n*:

- observed read depth and read pair information (*d_i_, p_i_*), i.e. *d_i_* is the total observed read depth, and *p_i_* is the number of discordantly mapped read pairs.
- potential duplicated or deleted or inverted region (*α_i_, β_i_*).
- potential breakpoint *γ_i_*.
- potential SV type.

Assuming observed read depth and number of discordant read pairs follow a Poisson distribution, *λ >* 0,

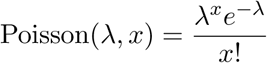

here, *λ* is the expected number of read depth or read pairs, and *x* is the observed number of read depth or read pairs respectively. However, the expected read depth or read pairs for some events might be zero, we approximate the probability by,

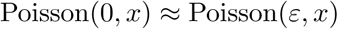

for a small *ε >* 0 (e.g. *ε* = 0.01 for read depth and *ε* = 0.001 for read pairs).

For each cluster *S_i_*, we define a random variable *state_i_* ∈ {0, 1, 2} in which the state of *S_i_* is *homozygous* if *state_i_* = 2, *heterozygous* if *state_i_* = 1, and *no event* if *state_i_* = 0. We also define a random variable *type_i_*, which represents the SV type for *S_i_*. Given *state_i_* = *k* and *type_i_* = *δ*, the likelihood of *S_i_* can be calculated as:

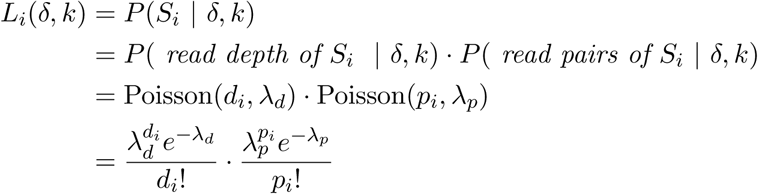

where *λ_d_* is the expected read depth of *S_i_* given *type_i_* = *δ, state_i_* = *k* and *λ_p_* is the expected read pairs of *S_i_* given *type_i_* = *δ, state_i_* = *k*.

We calculate *λ_d_* based on (*type_i_, state_i_*) and the expected read depth within the region (*α_i_, β_i_*) normalized with respect to its G+C content using a sliding window of size 100 bp, denoted by *E_d_*[(*α_i_, β_i_*)]. We calculate *λ_p_* based on the (*type_i_, state_i_*) and the expected number of discordantly mapped read pairs around the potential breakpoint *γ_i_*, denoted by *E_p_*[*γ_i_*]. For instance, if an event is categorized as homozygous deletion, we expect to see almost no read depth inside the potential deleted region (*α_i_, β_i_*), and the expected number of discordantly mapped read pairs should be approximately the expected number of reads containing the potential breakpoint, i.e *E_p_*[*γ_j_*]. For heterozygous deletion events, we expect to see half of the number of read depths and half of the expected number of discordantly mapped read pairs. We also calculate the likelihood score of no event at the potential region given that is categorized as deletion. For this case, we expect to see the expected number of read depths in that potential region and zero discordantly mapped read pairs. Similarly, the value for *λ_d_, λ_p_* can be approximately for inversion and duplications. Table 1 shows the value for *λ_d_, λ_p_* for each (*type_i_, state_i_*) using *E_d_*[(*α_i_, β_i_*)] and *E_p_*[*γ_i_*]. Note that even though the formulation for *λ_d_, λ_p_* are the same for all types of duplications, the likelihood score will be different because the potential regions (*α_i_, β_i_*) are different based on the categorized type of the event being considered. Furthermore, the read-pair support and signature will be different for each type of duplication which is the key in resolving the type of duplication.

**Table 1:** Formulation for *λ_d_* and *λ_p_* for maximum valid cluster *S_i_*

| SV Type | State | $\lambda_d$ | $\lambda_p$ |
| --- | --- | --- | --- |
| Deletion | *homozygous* | 0.01 | $E_p[\gamma_i]$ |
| | *heterozygous* | $0.5 \cdot E_d[(\alpha_i, \beta_i)]$ | $0.5 \cdot E_p[\gamma_i]$ |
| | *no event* | $E_d[(\alpha_i, \beta_i)]$ | 0.001 |
| Inversion | *homozygous* | $E_d[(\alpha_i, \beta_i)]$ | $E_p[\gamma_i]$ |
| | *heterozygous* | $E_d[(\alpha_i, \beta_i)]$ | $0.5 \cdot E_p[\gamma_i]$ |
| | *no event* | $E_d[(\alpha_i, \beta_i)]$ | 0.001 |
| Inverted Duplication | *homozygous* | $2 \cdot E_d[(\alpha_i, \beta_i)]$ | $E_p[\gamma_i]$ |
| | *heterozygous* | $1.5 \cdot E_d[(\alpha_i, \beta_i)]$ | $0.5 \cdot E_p[\gamma_i]$ |
| | *no event* | $E_d[(\alpha_i, \beta_i)]$ | 0.001 |
| Direct Duplication | *homozygous* | $2 \cdot E_d[(\alpha_i, \beta_i)]$ | $E_p[\gamma_i]$ |
| | *heterozygous* | $1.5 \cdot E_d[(\alpha_i, \beta_i)]$ | $0.5 \cdot E_p[\gamma_i]$ |
| | *no event* | $E_d[(\alpha_i, \beta_i)]$ | 0.001 |
| Tandem Duplication | *homozygous* | $2 \cdot E_d[(\alpha_i, \beta_i)]$ | $E_p[\gamma_i]$ |
| | *heterozygous* | $1.5 \cdot E_d[(\alpha_i, \beta_i)]$ | $0.5 \cdot E_p[\gamma_i]$ |
| | *no event* | $E_d[(\alpha_i, \beta_i)]$ | 0.001 |

#### 2.3.2 SV weight

For each potential SV we calculate a score to represent how likely a SV prediction is correct given the observed signature. Note that, for each SV, we calculate the likelihood considering homozygous state and heterozygous state separately.

We define the score as ratio of log of likelihoods of the putative SV being true given the observed data over it being false. Note that we use log function to avoid numerical errors. Even those the standard approach is to use logarithm of the ratio, we heuristically use the ratio to make sure that the scores are positive, which will work better for the set cover approximation algorithm we will use in the next step.

The score of potential SV *S_i_* is defined as follows:

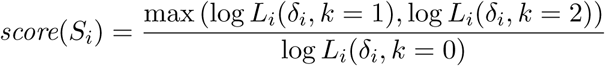

where *δ_i_* is the potential SV type of *S_i_*. Again, *k* = 0, 1, 2 implies that the state of *S_i_* is *no event*, *heterozygous*, *homozygous* respectively.

#### 2.3.3 Multi-mapping reads

We have previously showed that a greedy approach motivated by weighted-set cover problem performs well in discovery of SVs with multiple mapping of the reads [12]. It guaranties an *O*(log(*n*)) approximation. We therefore utilize a similar iterative greedy approach here as minimum weighted-set cover. More formally, at each step we select the set with the lowest ratio of SV score (*score*(*S_i_*)) and number of uncovered discordant paired end-reads being covered by that SV (*p_i_*)

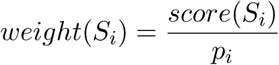

and continues this iterative process.

## 3 Results

### 3.1 Simulation

In order to evaluate performance of our SV detection algorithms, we generated a simulated genome first using VarSim [27]. VarSim “inserts” previously known real genomic variants into a given reference segment. Although it supports deletions, inversions, and tandem duplications, it does not yet simulate interspersed segmental duplications. Therefore we developed a new simulator called CNVSim to additionally simulate interspersed duplications in both direct and inverted duplication.

In total, we simulated SVs of lengths selected uniformly random between 500 bp and 10 Kbp. For inverted duplications and interspersed direct duplications, the distance from the new paralog to the original copy is chosen uniformly random between 5,000 bp and 50 Kbp. All segments are sampled randomly from the well-defined (i.e., no assembly gaps) regions in the reference genome, and guaranteed to be non-overlapping. Each simulated SV can be in homozygous or heterozygous state.

Based on the human reference genome (GRCh37), we simulated total of 1,200 SVs including 700 deletions, 579 inversions, 200 tandem duplications, 200 inverted duplications, and 200 interspersed direct duplications. We then simulated WGS data at four depth of coverages 10×, 20×, 30×, 60× using wgsim (https://github.com/lh3/wgsim). We mapped the reads back to the human reference genome (GRCh37) using BWA-MEM [21]. Finally we obtained structural variation call sets using TARDIS, DELLY [30], LUMPY [19], TIDDIT [11], and SoftSV [4].

We included analysis of all types of SVs in our simulation and real data experiments following our motivation we outlined in section 2.1 and Figures 1 and 2. We would like to reiterate that inability to call interspersed segmental duplications results in higher false positives in both deletion and inversion discovery. Through characterization of segmental duplications and integration of a read depth based probabilistic model, TARDIS achieves better inversion and deletion discovery accuracy by correct classification of more complex SV types. Further analysis on the simulations revealed that 95 of 773 deletions predicted by LUMPY and 96 of 852 deletions predicted by DELLY are indeed interspersed duplications in direct orientation. Similarly, 109 of 1,286 DELLY-predicted inversions were in fact inverted segmental duplications.

Finally, we simulated 10 large (up to 1 Mbp) segmental duplications in chromosome Y to assess the power of TARDIS in detecting large duplications. TARDIS correctly identified 4/10 duplications of size >63 Kb (Supplementary Table S1).

Table 2 shows the true positive rate (TPR) and false discovery rate (FDR) of TARDIS compared to DELLY, LUMPY, TIDDIT and SoftSV on the simulated data. TARDIS achieved a substantially higher TPR and a lower FDR for deletions and duplications overall. Additionally, its sensitivity is comparable to LUMPY and SoftSV in terms of inversion predictions. (See Supplementary Figure 2 for precision-recall curves of inversions and duplications.)

**Table 2:**
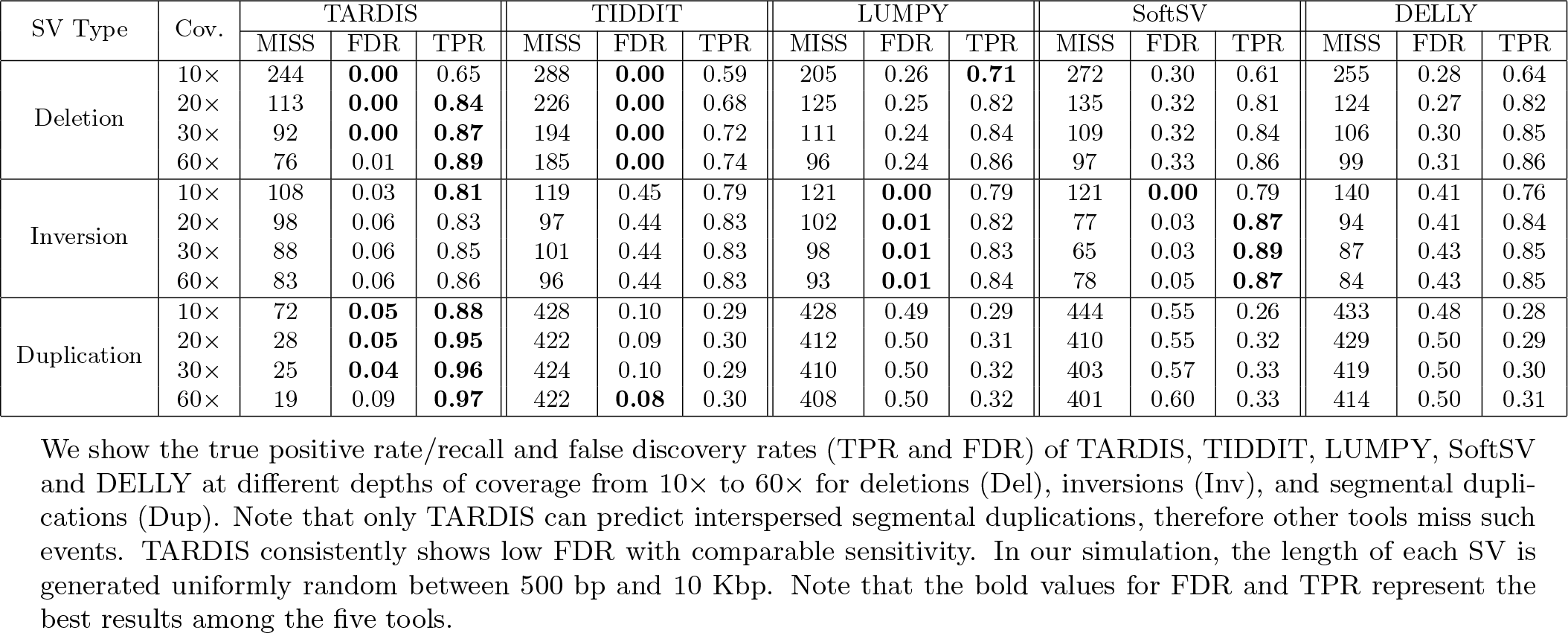
Summary of simulation predictions by TARDIS, TIDDIT, LUMPY, SoftSV and DELLY.

| SV Type | Cov. | TARDIS |  |  | TIDDIT |  |  | LUMPY |  |  | SoftSV |  |  | DELLY |  |  |
| --- | --- | --- | --- | --- | --- | --- | --- | --- | --- | --- | --- | --- | --- | --- | --- | --- |
|  |  | MISS | FDR | TPR | MISS | FDR | TPR | MISS | FDR | TPR | MISS | FDR | TPR | MISS | FDR | TPR |
| Deletion | 10× | 244 | **0.00** | 0.65 | 288 | **0.00** | 0.59 | 205 | 0.26 | **0.71** | 272 | 0.30 | 0.61 | 255 | 0.28 | 0.64 |
|  | 20× | 113 | **0.00** | **0.84** | 226 | **0.00** | 0.68 | 125 | 0.25 | 0.82 | 135 | 0.32 | 0.81 | 124 | 0.27 | 0.82 |
|  | 30× | 92 | **0.00** | **0.87** | 194 | **0.00** | 0.72 | 111 | 0.24 | 0.84 | 109 | 0.32 | 0.84 | 106 | 0.30 | 0.85 |
|  | 60× | 76 | 0.01 | **0.89** | 185 | **0.00** | 0.74 | 96 | 0.24 | 0.86 | 97 | 0.33 | 0.86 | 99 | 0.31 | 0.86 |
| Inversion | 10× | 108 | 0.03 | **0.81** | 119 | 0.45 | 0.79 | 121 | **0.00** | 0.79 | 121 | **0.00** | 0.79 | 140 | 0.41 | 0.76 |
|  | 20× | 98 | 0.06 | 0.83 | 97 | 0.44 | 0.83 | 102 | **0.01** | 0.82 | 77 | 0.03 | **0.87** | 94 | 0.41 | 0.84 |
|  | 30× | 88 | 0.06 | 0.85 | 101 | 0.44 | 0.83 | 98 | **0.01** | 0.83 | 65 | 0.03 | **0.89** | 87 | 0.43 | 0.85 |
|  | 60× | 83 | 0.06 | 0.86 | 96 | 0.44 | 0.83 | 93 | **0.01** | 0.84 | 78 | 0.05 | **0.87** | 84 | 0.43 | 0.85 |
| Duplication | 10× | 72 | **0.05** | **0.88** | 428 | 0.10 | 0.29 | 428 | 0.49 | 0.29 | 444 | 0.55 | 0.26 | 433 | 0.48 | 0.28 |
|  | 20× | 28 | **0.05** | **0.95** | 422 | 0.09 | 0.30 | 412 | 0.50 | 0.31 | 410 | 0.55 | 0.32 | 429 | 0.50 | 0.29 |
|  | 30× | 25 | **0.04** | **0.96** | 424 | 0.10 | 0.29 | 410 | 0.50 | 0.32 | 403 | 0.57 | 0.33 | 419 | 0.50 | 0.30 |
|  | 60× | 19 | 0.09 | **0.97** | 422 | **0.08** | 0.30 | 408 | 0.50 | 0.32 | 401 | 0.60 | 0.33 | 414 | 0.50 | 0.31 |

We show the true positive rate/recall and false discovery rates (TPR and FDR) of TARDIS, TIDDIT, LUMPY, SoftSV and DELLY at different depths of coverage from 10× to 60× for deletions (Del), inversions (Inv), and segmental duplications (Dup). Note that only TARDIS can predict interspersed segmental duplications, therefore other tools miss such events. TARDIS consistently shows low FDR with comparable sensitivity. In our simulation, the length of each SV is generated uniformly random between 500 bp and 10 Kbp. Note that the bold values for FDR and TPR represent the best results among the five tools.

In these simulation experiments we used the default variables, which require at least 5 read pairs that support the SV event. Although this cut off works well, it contributes to higher number of false positives when the depth of coverage is high (Table 2). To demonstrate the effects of the values for this parameter, we repeated the experiment with varying minimum number of read pair support values. We confirmed that with higher values, we can reduce the FDR for high coverage genomes (Supplementary Table S2).

Furthermore, TARDIS can classify duplications into tandem, interspersed directed duplication and inverted duplication. However, DELLY, LUMPY, TIDDIT and SoftSV are not designed to characterize interspersed segmental duplications, therefore we cannot provide comparisons. Table 3 shows the TDR, FDR, and the exact count of the number of True/False predictions for each type of segmental duplication.

**Table 3:** Characterization of different types of segmental duplications using TARDIS on simulated data.

| Duplication Type | Coverage | # SVs | Missed | True | TPR | False | FDR |
| --- | --- | --- | --- | --- | --- | --- | --- |
| Inverted Interspersed Duplication | 10× | 200 | 15 | 185 | 0.93 | 7 | 0.04 |
|  | 20× | 200 | 10 | 190 | 0.95 | 11 | 0.05 |
|  | 30× | 200 | 12 | 188 | 0.94 | 15 | 0.07 |
|  | 60× | 200 | 9 | 191 | 0.96 | 33 | 0.15 |
| Direct Interspersed Duplication | 10× | 200 | 10 | 190 | 0.95 | 3 | 0.02 |
|  | 20× | 200 | 7 | 193 | 0.97 | 0 | 0.00 |
|  | 30× | 200 | 6 | 194 | 0.97 | 4 | 0.02 |
|  | 60× | 200 | 5 | 195 | 0.98 | 9 | 0.04 |
| Tandem Duplication | 10× | 200 | 47 | 153 | 0.77 | 21 | 0.12 |
|  | 20× | 200 | 11 | 189 | 0.95 | 15 | 0.07 |
|  | 30× | 200 | 7 | 193 | 0.97 | 10 | 0.05 |
|  | 60× | 200 | 5 | 195 | 0.98 | 16 | 0.08 |

TARDIS can classify duplications into tandem, interspersed directed duplication and inverted duplication. However, DELLY, LUMPY, TIDDIT and SoftSV are not designed to characterize these complex SVs. This table shows the true positive rate (recall) and false discovery rate (TPR and FDR respectively) of TARDIS for each type of duplication.

### 3.2 Haploid genome analyses

As the first experiment with real data sets, we downloaded short read HTS data generated from two haploid cell lines, namely CHM1 and CHM13 [16, 38]. We mapped the reads to human reference genome (GRCh37) using BWA-MEM [21]. We also obtained call sets generated with PacBio data from the same genomes [6], but here we use updated SV calls (Mark Chaisson, personal communication), which we use as the true inversion set to compare with our predictions.

We present the comparison of the inversion predictions made by TARDIS and two state of the art methods LUMPY and DELLY in Figure 4. Note that we only consider inversions of length > 100 bp. Figure 4) (a) & (b) show the comparison of TARDIS predictions with those of other tools on CHM1 and CHM13 respectively (We also present a similar comparison for deletion predictions in Supplementary Figure 3). Overall, TARDIS achieves better accuracy. We also tested the highest scoring set (n=50) of predicted inversions by each tool generated for the CHM1 genome. Briefly, we used a reference-guided *de novo* assembly of PacBio reads generated from the same genome [6] and mapped the contigs to the loci of interest. We show a receiver-operating-charasteristic-like plot that uses actual numbers of true and false calls instead of rates (TPR/FDR) (Supplementary Figure 4). Here we observe that compared to LUMPY and DELLY, TARDIS achieves better area under the curve. However, we note that the main reason for DELLY and LUMPY curves being closer to that of TARDIS for low number of false calls is because there were several predictions for which corresponding contigs did not exist in the assembled genome, therefore omitted from this plot.

**Figure 4:**
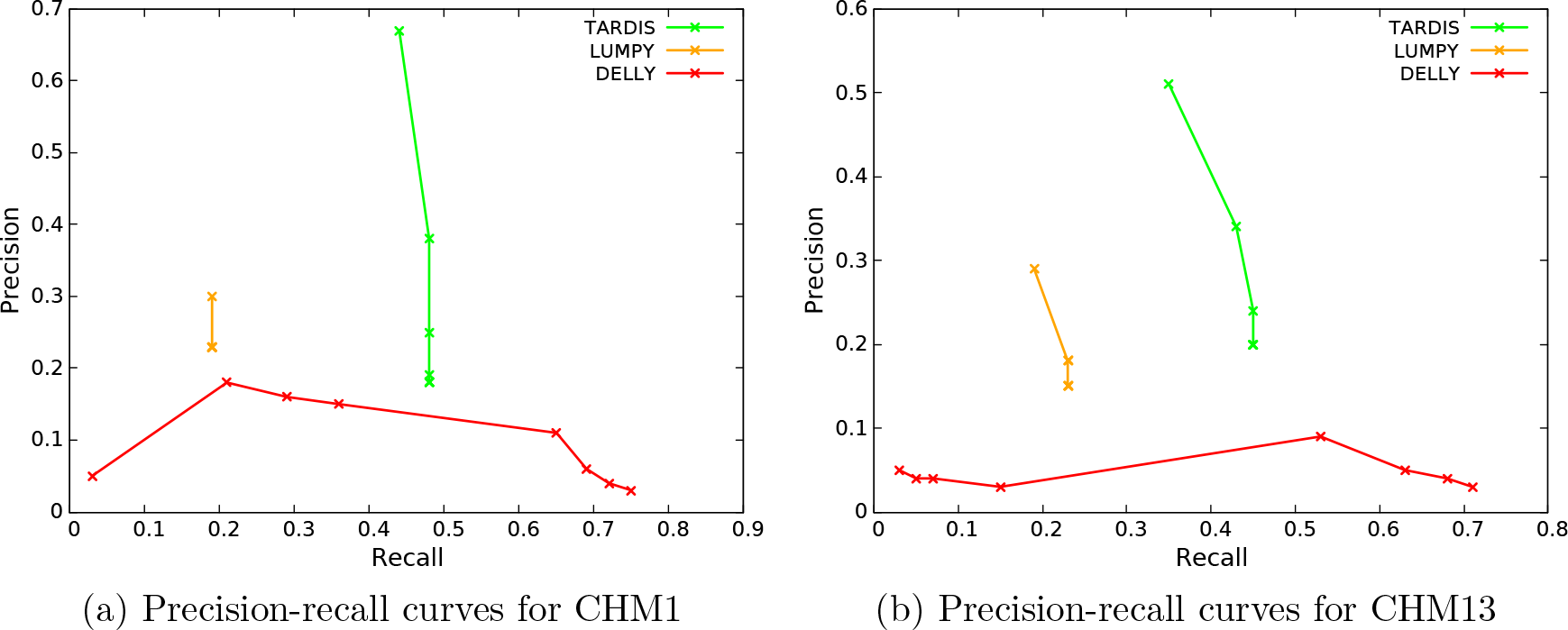
Precision - Recall curves for the comparison of inversion predictions on CHM1 and CHM13 genomes. Overall TARDIS achieves better accuracy than the two other approaches tested. (a), (b) comparison of CHM1 and CHM13 predicted inversions using PacBio reads based on BLASR mappings.

We provide the full set of the 50 highest scoring segmental duplications that TARDIS predicts in the CHM1 genome together with *in silico* validation using the corresponding PacBio-based assembly (Table 4). Almost all of the predicted duplications, except one, were validated using long reads. We provide the PacBio alignments of some of these events and top 20 highest scoring CHM13 predictions in Supplementary Materials. Note that in most cases TARDIS assigned the correct subtype of duplications (inverted, direct or tandem duplication) to the prediction. As expected, the highest number of segmental duplications in the top 50 were tandem duplications (> 50% of all duplications).

**Table 4:** 50 highest scoring segmental duplications predicted by TARDIS in the CHM1 genome.

| Duplication<br>Insertion Locus |  |  | TARDIS |  | Validation<br>(PacBio) | Duplication<br>Insertion Locus |  |  | TARDIS |  | Validation<br>(PacBio) |
| --- | --- | --- | --- | --- | --- | --- | --- | --- | --- | --- | --- |
|  |  |  | Dup. Type | Score |  |  |  |  | Dup. Type | Score |  |
| chr11 | 63,701,552 | - 63,702,044 | Direct | 0.000096 | True | chr2 | 37,928,294 | - 38,101,823 | Tandem | 0.000073 | N/A |
| chr3 | 194,546,158 | - 194,546,552 | Direct | 0.000100 | True | chr20 | 60,032,848 | - 60,033,403 | Tandem | 0.000080 | True |
| chr5 | 143,512,369 | - 143,512,967 | Direct | 0.000139 | True | chr5 | 3,323,855 | - 3,324,309 | Tandem | 0.000106 | N/A |
| chr2 | 240,640,651 | - 240,641,122 | Direct | 0.000199 | True | chr7 | 2,554,464 | - 2,554,791 | Tandem | 0.000111 | True |
| chr20 | 2,359,605 | - 2,360,003 | Direct | 0.000271 | True | chr12 | 110,099,340 | - 110,099,746 | Tandem | 0.000117 | True |
| chr9 | 112,285,747 | - 112,286,145 | Direct | 0.000300 | True | chr6 | 168,052,194 | - 168,052,468 | Tandem | 0.000117 | True |
| chr8 | 2,215,143 | - 2,215,392 | Direct | 0.000310 | N/A | chr1 | 207,097,489 | - 207,097,910 | Tandem | 0.000123 | True |
| chr18 | 69,711,702 | - 69,712,115 | Direct | 0.000323 | True | chr16 | 86,008,734 | - 86,009,147 | Tandem | 0.000127 | True |
| chr17 | 46,615,512 | - 46,615,903 | Direct | 0.000326 | True | chr17 | 80,317,607 | - 80,318,019 | Tandem | 0.000127 | N/A |
| chr6 | 160,877,582 | - 160,878,047 | Direct | 0.000342 | N/A | chr10 | 127,513,435 | - 127,513,672 | Tandem | 0.000129 | True |
| chr2 | 125,052,915 | - 125,053,261 | Inverted | 0.000088 | True | chr14 | 106,049,125 | - 106,049,349 | Tandem | 0.000129 | True |
| chr3 | 43,834,996 | - 43,835,748 | Inverted | 0.000089 | True | chr6 | 44,012,338 | - 44,012,957 | Tandem | 0.000129 | True |
| chr14 | 67,171,710 | - 67,172,020 | Inverted | 0.000092 | True | chr9 | 132,158,817 | - 132,159,088 | Tandem | 0.000129 | N/A |
| chr2 | 72,440,071 | - 72,440,597 | Inverted | 0.000105 | True | chr12 | 13,164,470 | - 13,164,800 | Tandem | 0.000136 | True |
| chr9 | 107,816,537 | - 107,817,079 | Inverted | 0.000140 | True | chr20 | 62,720,020 | - 62,720,215 | Tandem | 0.000136 | True |
| chr17 | 36,405,748 | - 36,407,397 | Inverted | 0.000149 | False | chr10 | 132,974,754 | - 132,975,320 | Tandem | 0.000144 | True |
| chr1 | 114,645,858 | - 114,646,155 | Inverted | 0.000235 | True | chr8 | 2,215,817 | - 2,216,236 | Tandem | 0.000144 | N/A |
| chr5 | 115,350,905 | - 115,351,086 | Inverted | 0.000236 | True | chr9 | 34,681,581 | - 34,681,899 | Tandem | 0.000194 | True |
| chr12 | 71,532,699 | - 71,533,378 | Inverted | 0.000245 | True | chr6 | 35,754,661 | - 35,766,731 | Tandem | 0.000255 | True |
| chr7 | 31,586,861 | - 31,587,129 | Inverted | 0.000278 | True | chr20 | 62,123,612 | - 62,124,210 | Tandem | 0.000257 | True |
| chr18 | 11,511,287 | - 11,511,480 | Inverted | 0.000280 | True | chr20 | 59,567,884 | - 59,590,251 | Tandem | 0.000268 | True |
|  |  |  |  |  |  | chr18 | 77,831,329 | - 77,831,784 | Tandem | 0.000273 | N/A |
|  |  |  |  |  |  | chrX | 417,958 | - 418,361 | Tandem | 0.000273 | True |
|  |  |  |  |  |  | chr20 | 42,325,214 | - 42,325,573 | Tandem | 0.000290 | True |
|  |  |  |  |  |  | chr19 | 34,882,471 | - 34,883,258 | Tandem | 0.000310 | True |
|  |  |  |  |  |  | chr2 | 3,184,299 | - 3,185,046 | Tandem | 0.000310 | N/A |
|  |  |  |  |  |  | chr3 | 197,117,159 | - 197,117,807 | Tandem | 0.000318 | N/A |

Here we list the insertion locations of the top 50 scoring segmental duplications in CHM1 genome. All predictions are sorted by the SV score (lower is better). If the validation is N/A, that means the incorrect prediction from PacBio data, which will be skipped in the comparison. TARDIS only gives one false call and three interspersed duplications that are wrongly assigned to tandem duplications.

### 3.3 NA12878 genome

We also analyzed the WGS data generated from NA12878 using TARDIS for various types of SV discovery and compared the results against state-of-the-art methods for inversion prediction. Similar to the simulation and CHM1/13 results, TARDIS outperformed the tested methods for SV discovery (see Supplementary Figure 16 for inversion comparison with a set of validated inversions on this sample).

More interestingly, we have found an example of a large inverted duplication in NA12878 sample which we validated using available orthogonal PacBio data generated from the same sample (Figure 5). The interesting point about this inverted duplication is that it is larger than 10 Kbp and the distance between locus of insertion and the duplicated region is also larger, which shows a potential start of a new segmental duplication.

**Figure 5:**
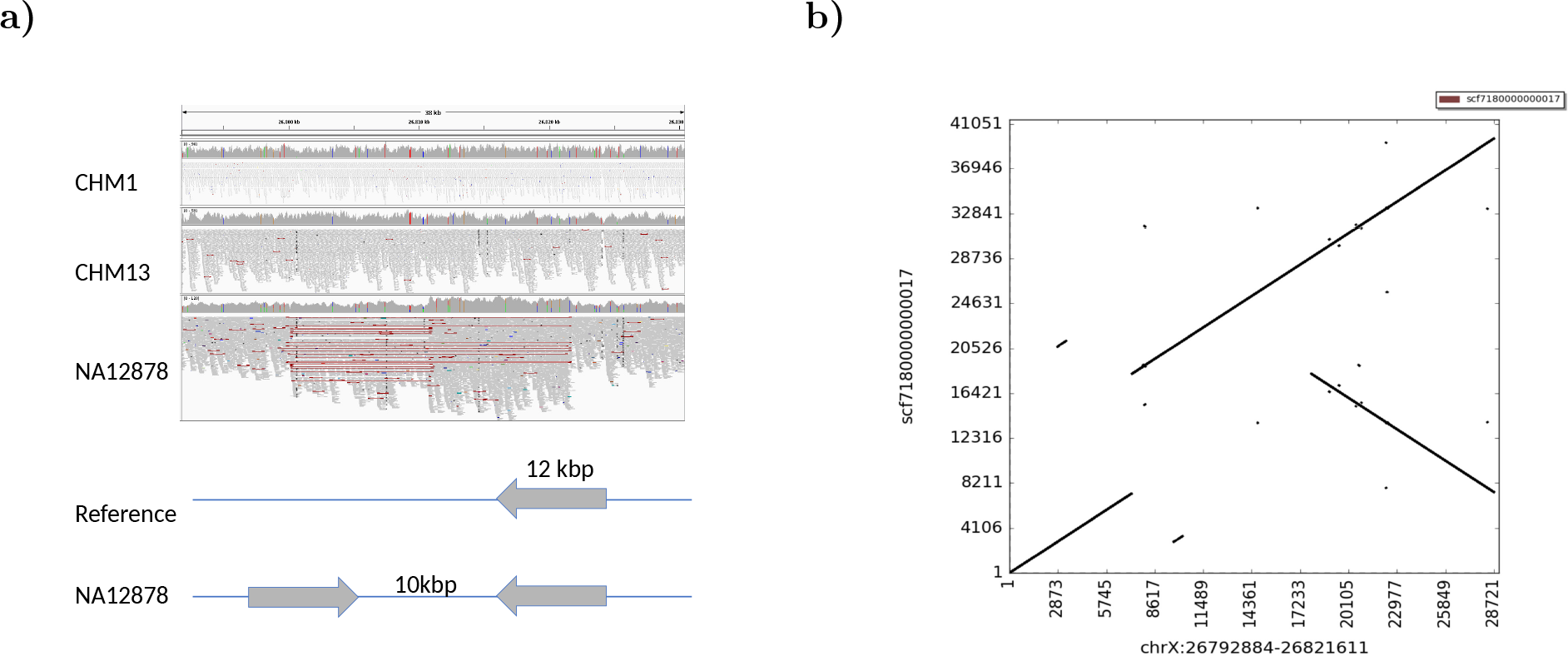
a) Illumina read mapping information visualized using IGV[42]. Here the read pairs in the NA12878 genome show typical inversion signature (red lines), where all reads map concordantly in CHM1 and CHM13 genomes, and a simple sketch of the alternative inverted duplication structure of the same region. b) Dot plot matrix validation using PacBio data, which shows an inverted duplication. The whole genome assembly shows an inverted duplication of a 12 Kb segment separated by 10 Kb. This region demonstrates the case where read pair based clustering confuses an inverted duplication with a simple inversion.

## 4 Discussion

Characterization of structural variants using HTS data is a well-studied problem. Still, due to the difficulty of accurately predicting complex variants, most of the current approaches mainly focus on specific forms of SVs. In this paper we describe novel algorithms to detect complex SV events such as tandem, direct and inverted interspersed segmental duplications simultaneously with simpler forms SV using whole genome sequencing data. Our approach integrates multiple sequence signatures to identify and cluster potential SV regions under the assumption of maximum parsimony. However, complex SV events usually generate similar signatures (i.e., inversion vs. inverted duplication), which make it difficult to differentiate particular SV types. Therefore, we strengthened our method by using a probabilistic likelihood model to overcome this obstacle by calculating a likelihood score for each SV.

Using simulated and real data sets, we showed that TARDIS outperforms state-of-the art methods in terms of specificity for all types of SVs, and achieves considerably high true discovery rate for segmental duplications with moderate time and memory requirements (See Supplementary Table S4 for a comparison of different tools for CHM1 and NA12878 genomes.). It should be noted that it TARDIS is currently one of the few methods that can classify duplications as tandem and interspersed in direct or inverted orientation using HTS data. Additionally, it demonstrates comparable sensitivity in deletions and inversions.

Here we only focused on tandem duplications in direct orientation, although inverted tandem repeats in genomes, or DNA palindromes, also exist especially in the human Y chromosome [5, 43]. However, these DNA palindromes were incorporated in the human genome over millions of years of evolution, and polymorphic inverted tandem duplication events are rare. Because of this, the mechanisms forming DNA palindromes are not yet well-established and we are not aware of a resource of validated DNA palindrome polymorphisms. We therefore ignore such variants in this study and we aim to address them in the future.

Future improvements in TARDIS will include addition of local assembly signature to help it achieve better accuracy. Although simulation experiments demonstrated potential efficacy of TARDIS in segmental duplication predictions, those that are generated from real genomes need to be experimentally verified to fully understand the power and shortcomings of the TARDIS algorithm. We can then apply TARDIS to thousands of genomes that were already sequenced as part of various projects, such as the 1000 Genomes Project to advance our understanding of the SV spectrum in human genomes. Another possible direction for TARDIS can be integration of new methods to better detect somatic structural variation detection, which we can then apply to cancer genomes.

## Acknowledgements

We thank E. Ebren and F. Karaoglanoglu for their help in creating simulation data sets. We would thank Evan E. Eichler for insightful advise and comments. Part of the work was done during FH postdoc training in Evan E. Eichler’s lab. We would also like to Mark Chaisson for providing PacBio call sets for CHM1 and CHM13, and the local assembly of these genomes.

## Funding

This work was supported by a grant by a TÜBİTAK (215E172), and an EMBO Installation Grant (IG-2521) to C.A., and an NSF grant (1528234) to F.H. The authors also acknowledge the Computational Genomics Summer Institute funded by NIH grant GM112625 that fostered international collaboration among the groups involved in this project.

## Availability

TARDIS is available under BSD 3-clause license at https://github.com/BilkentCompGen/tardis, and the CNVSim simulator is available at https://github.com/LeMinhThong/CNVSim. NA12878 WGS data set can be downloaded from https://www.illumina.com/platinumgenomes.html. SRA IDs for CHM1 and CHM13 are SRP044331 and SRP080317, respectively. GenBank assembly accession numbers for CHM1 and CHM13 assemblies are GCA_000306695.2 and GCA_000983455.2.

