## Supplementary Material for "Discovery of tandem and interspersed segmental duplications using high throughput sequencing"

### 4.1 Command lines

#### 4.1.1 Simulation using VarSim

In order to simulate SVs including deletions, inversions and tandem duplications we used VarSim, which inserts known genomic variants into a given reference genome. However, it is unable to simulate interspersed duplications, thus we developed a new simulator called CNVSim to include interspersed duplications in direct and inverted orientations to the simulated genome. We additionally added some fixed real inversions to make the simulation more realistic. Finally, we created a VCF file including the interspersed duplications and some of the real inversions and used it as input to VarSim to generate the fastq files encompassing all the genomic variants.

```
vc_in_vcf=/share/varsim_files/All.vcf.gz
sv_insert_seq=/share/varsim_files/insert_seq.txt
sv_dgv=/share/varsim_files/GRCh37_hg19_supportingvariants_2013-07-23.txt
reference=human_g1k_v37_gatk.fasta
simulator_executable=/share/varsim_files/ART/art_bin_VanillaIceCream/art_illumina
vcf=invdup_simu.vcf

varsim.sh --reference $reference --id simu --read_length 100 --sv_num_ins 0 --sv_num_del 5[illegible]
--sv_num_dup 500 --sv_num_inv 500 --sv_percent_novel 0.01  --mean_fragment_size 350 \
--sd_fragment_size 50 --sv_min_length_lim 50 --sv_max_length_lim 10000 --sv_insert_seq $sv[illegible]
--vc_in_vcf $vc_in_vcf --sv_dgv $sv_dgv --nlanes 1 --total_coverage $coverage \
--simulator_executable $simulator_executable --out_dir $out --log_dir $log --work_dir $wor[illegible]
--simulator art --vcfs $vcf
```

#### 4.1.2 SV Discovery Tools

**TIDDIT**:

```
python TIDDIT.py --sv --bam CHM1.bam --ref human_g1k_v37_gatk.fasta -o chm1
```

**LUMPY**:

```
lumpyexpress -B CHM1.bam -o chm1.vcf
```

**DELLY**:

```
delly call -o chm1 -g human_g1k_v37_gatk.fasta -x excludeTemplates/human.hg19.excl.tsv CHM[illegible]
```

**SoftSV**:

```
SoftSV --input CHM1.bam --output chm1
```

**TARDIS**:

```
tardis -i CHM1.bam --ref human_g1k_v37_gatk.fasta --sonic human_g1k_v37.sonic --out chm1
```

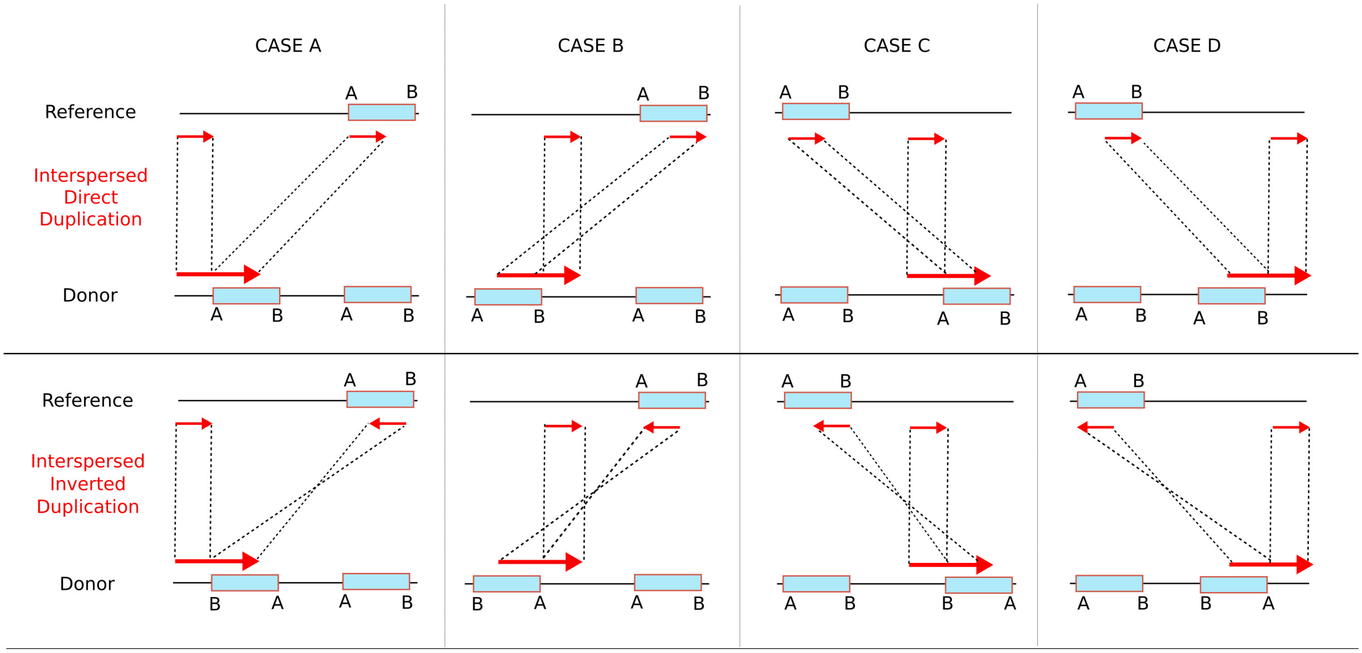


Supplementary Fig. 1: Split Read signatures for inverted and direct interspersed segmental duplications. In case A and B, duplicated part is inserted on the left and in C and D, on the right of the original segment. A) Soft clip is at the end of the read and is mapped after the primary mapping. B) Soft clip is at the beginning of the read and is mapped after the primary mapping. C) Soft clip is at the end of the read and is mapped before the primary mapping. D) Soft clip is at the beginning of the read and is mapped before the primary mapping.

Supplementary Table S1: Large segmental duplications found in chromosome Y simulation.

| **chromosome** | **start** | **end** | **length** | **type** | **genotype** | **distance to insertio[illegible]** |
|---|---|---|---|---|---|---|
| Y | 5,580,120 | 5,648,940 | 68,820 | direct interspersed | homozygous | [illegible] |
| Y | 15,349,440 | 15,442,800 | 93,360 | tandem | homozygous | |
| Y | 17,107,980 | 17,171,160 | 63,180 | tandem | homozygous | |
| Y | 18,553,380 | 18,670,200 | 116,820 | tandem | heterozygous | |

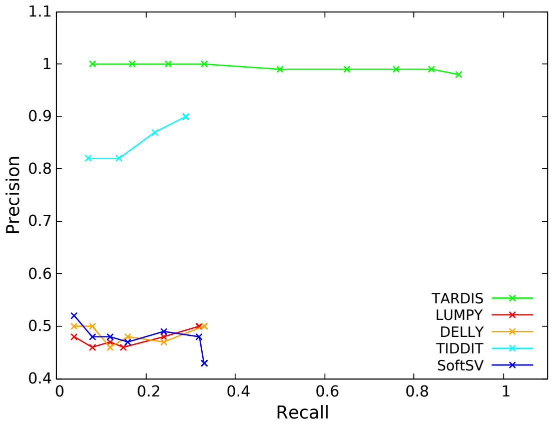


(a) Precision-recall curves for duplication predictions

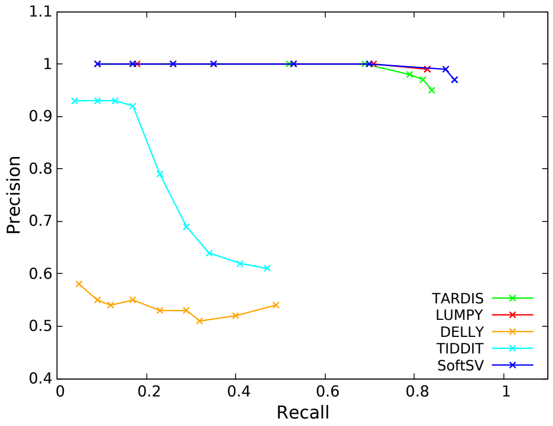


(b) Precision-recall curves for inversion predictions

Supplementary Fig. 2: Precision-Recall curves for the comparison of duplication (a) and inversion (b) predictions on the simulated dataset for 30X coverage using TARDIS, TIDDIT, LUMPY, DELLY and SoftSV.

Supplementary Table S2: Effect of Read-Pair Support for SV discovery for TARDIS

| RP support cut-off | SV Type | # SVs | True | False | Miss | FDR | TPR |
|---|---|---|---|---|---|---|---|
| 5 | Deletion | 700 | 624 | 5 | 76 | 0.01 | 0.89 |
| | Inversion | 579 | 496 | 34 | 83 | 0.06 | 0.86 |
| | Interspersed Dups. | 200 | 195 | 9 | 5 | 0.04 | 0.98 |
| | Inverted Dups. | 200 | 191 | 33 | 9 | 0.15 | 0.96 |
| | Tandem Dups. | 200 | 194 | 26 | 6 | 0.12 | 0.97 |
| 8 | Deletion | 700 | 615 | 2 | 85 | 0.00 | 0.88 |
| | Inversion | 579 | 493 | 32 | 86 | 0.06 | 0.85 |
| | Interspersed Dups. | 200 | 195 | 17 | 5 | 0.08 | 0.98 |
| | Inverted Dups. | 200 | 191 | 28 | 9 | 0.13 | 0.96 |
| | Tandem Dups. | 200 | 191 | 22 | 9 | 0.10 | 0.96 |
| 9 | Deletion | 700 | 609 | 2 | 91 | 0.00 | 0.87 |
| | Inversion | 579 | 491 | 31 | 88 | 0.06 | 0.85 |
| | Interspersed Dups. | 200 | 194 | 8 | 6 | 0.04 | 0.97 |
| | Inverted Dups. | 200 | 190 | 28 | 9 | 0.13 | 0.96 |
| | Tandem Dups. | 200 | 191 | 20 | 9 | 0.09 | 0.96 |
| 10 | Deletion | 700 | 608 | 1 | 92 | 0.00 | 0.87 |
| | Inversion | 579 | 491 | 31 | 88 | 0.06 | 0.85 |
| | Interspersed Dups. | 200 | 194 | 8 | 6 | 0.04 | 0.97 |
| | Inverted Dups. | 200 | 190 | 27 | 10 | 0.12 | 0.95 |
| | Tandem Dups. | 200 | 190 | 20 | 10 | 0.10 | 0.95 |
| 20 | Deletion | 700 | 581 | 1 | 119 | 0.00 | 0.83 |
| | Inversion | 579 | 476 | 25 | 103 | 0.05 | 0.82 |
| | Interspersed Dups. | 200 | 193 | 5 | 7 | 0.03 | 0.97 |
| | Inverted Dups. | 200 | 188 | 20 | 12 | 0.10 | 0.94 |
| | Tandem Dups. | 200 | 176 | 20 | 24 | 0.10 | 0.88 |

Table shows the effect of read-pair support cut-off value in SV discovery accuracy, that is, the number of minimum supporting read-pairs for an SV to be selected. The analysis were done using the simulated data of 60x coverage. FDR and TPR denotes false discovery and true positive/recall rates respectively.

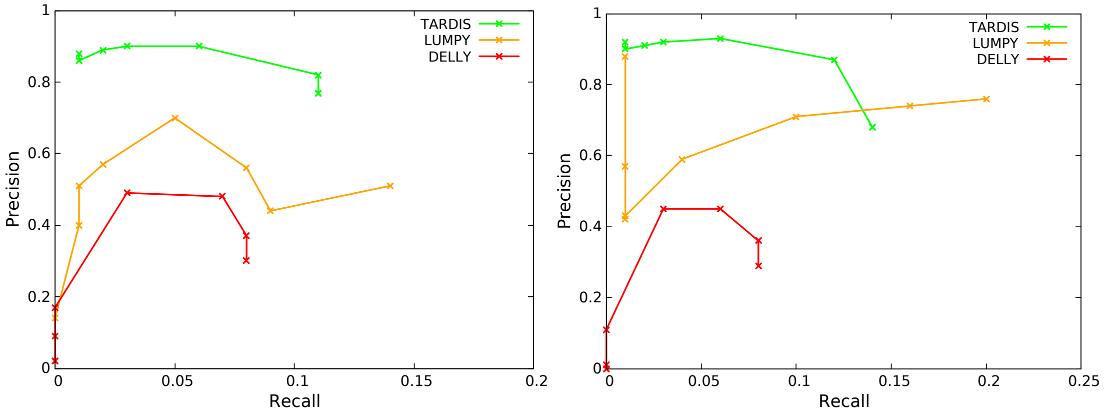


(a) Precision-recall curves for CHM1 (b) Precision-recall curves for CHM13

Supplementary Fig. 3: Precision - Recall curve for the comparison of deletion predictions on CHM1 and CHM13 genomes. Overall TARDIS achieves better accuracy than the two other approaches tested. (a), (b) comparison of CHM1 and CHM13 predicted inversions using PacBio reads based on BLASR mappings.

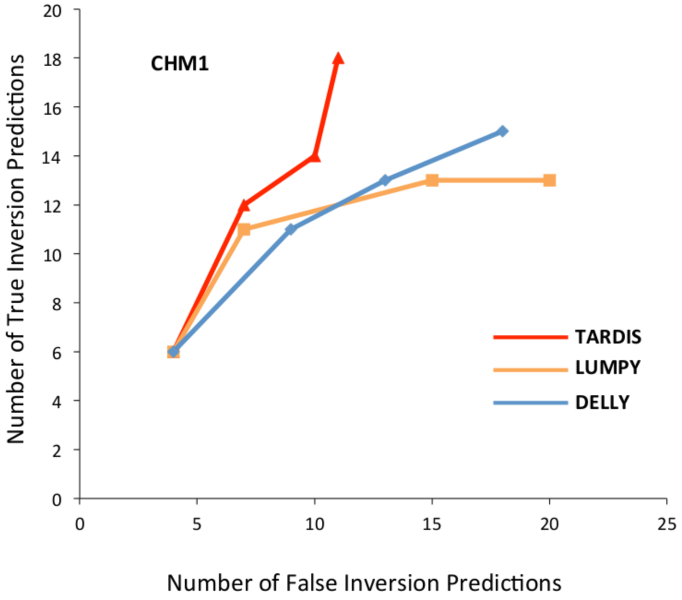


Supplementary Fig. 4: Validation of top predicted inversions of different tools using local assembly of the PacBio reads for CHM1.

Supplementary Table S3: 20 highest scoring segmental duplications predicted by TARDIS in the CHM13 genome.

| Duplication Insertion Locus | | | | TARDIS Dup. Type | TARDIS Score | Duplication Insertion Locus | | | | TARDIS Dup. Type | TARDIS Score |
|---|---|---|---|---|---|---|---|---|---|---|---|
| chr8 | 58,116,437 | - | 58,118,469 | Direct | 0.000188 | chr17 | 80,864,373 | - | 81,006,658 | Tandem | 0.000054 |
| chr6 | 119,011,599 | - | 119,012,388 | Direct | 0.000217 | chr16 | 81,798,809 | - | 81,799,175 | Tandem | 0.000121 |
| chr6 | 57,209,065 | - | 57,297,292 | Direct | 0.000233 | chr2 | 87,623,860 | - | 87,642,147 | Tandem | 0.000125 |
| chr5 | 143,512,394 | - | 143,512,967 | Direct | 0.000234 | chr19 | 34,882,364 | - | 34,882,984 | Tandem | 0.000216 |
| chr11 | 63,701,560 | - | 63,702,044 | Direct | 0.000234 | chr22 | 49,780,535 | - | 49,780,919 | Tandem | 0.000273 |
| chr8 | 76,769,879 | - | 76,770,323 | Direct | 0.000247 | chr5 | 1,044,880 | - | 1,045,357 | Tandem | 0.000273 |
| chr3 | 194,546,160 | - | 194,546,552 | Direct | 0.000261 | chr6 | 44,012,353 | - | 44,012,977 | Tandem | 0.000273 |
| chr5 | 140,859,762 | - | 140,860,171 | Direct | 0.000269 | | | | | | |
| chr14 | 48,325,266 | - | 48,325,533 | Inverted | 0.000208 | | | | | | |
| chr11 | 98,844,907 | - | 98,845,328 | Inverted | 0.000210 | | | | | | |
| chr2 | 61,703,139 | - | 61,703,479 | Inverted | 0.000210 | | | | | | |
| chr5 | 169,597,408 | - | 169,597,797 | Inverted | 0.000220 | | | | | | |
| chr12 | 78,387,851 | - | 78,388,292 | Inverted | 0.000246 | | | | | | |

Here we list the insertion locations of the top 20 scoring segmental duplications in CHM13 genome. All predictions are sorted by the SV score (lower is better).

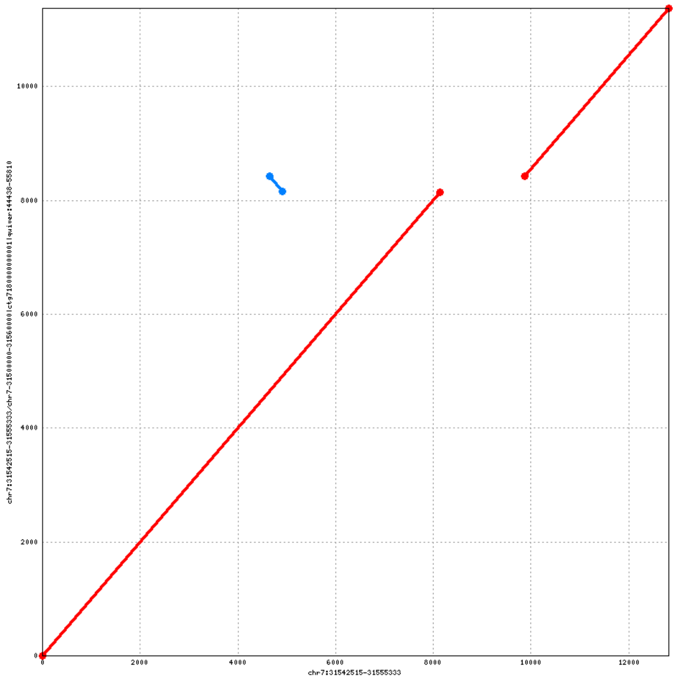


Supplementary Fig. 5: Inversion predicted within 7:31,586,823-31,590394.

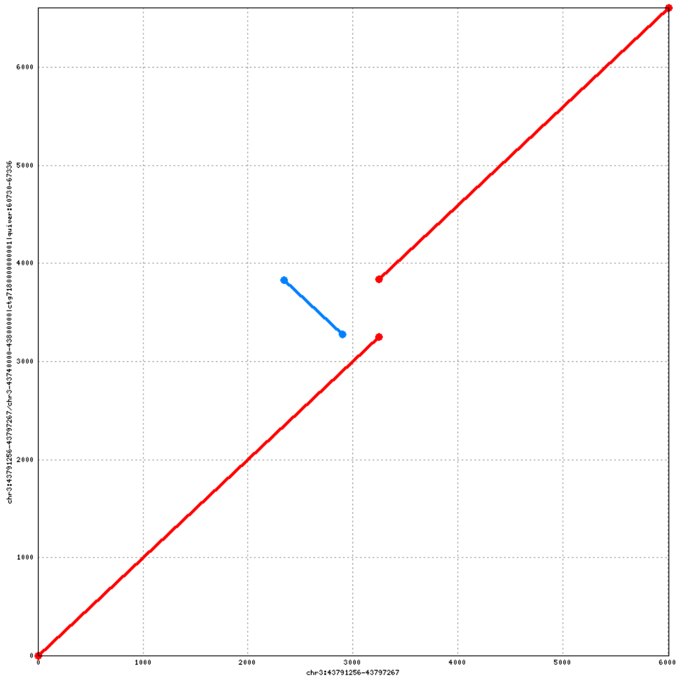


Supplementary Fig. 6: Inverted Duplication predicted within 3:43,834,994-43,836,299.

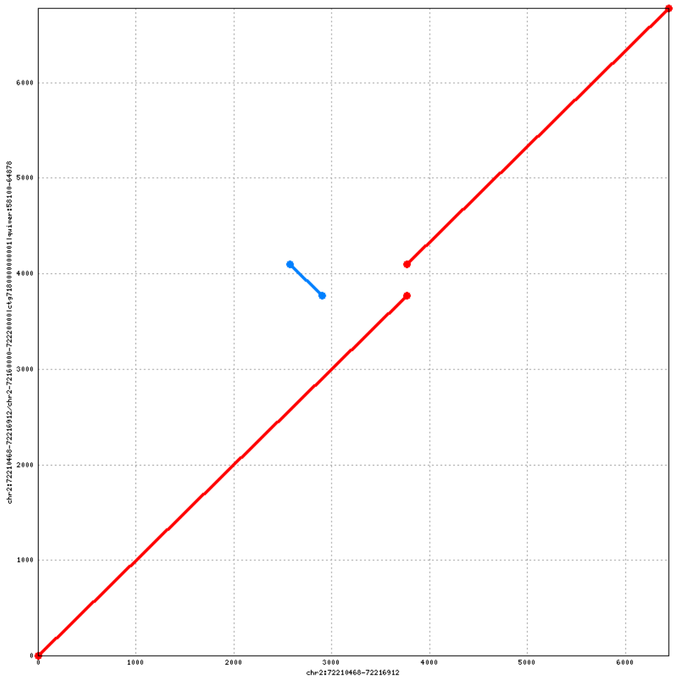


Supplementary Fig. 7: Inverted Duplication predicted within 2:72,440,066-72,441,647.

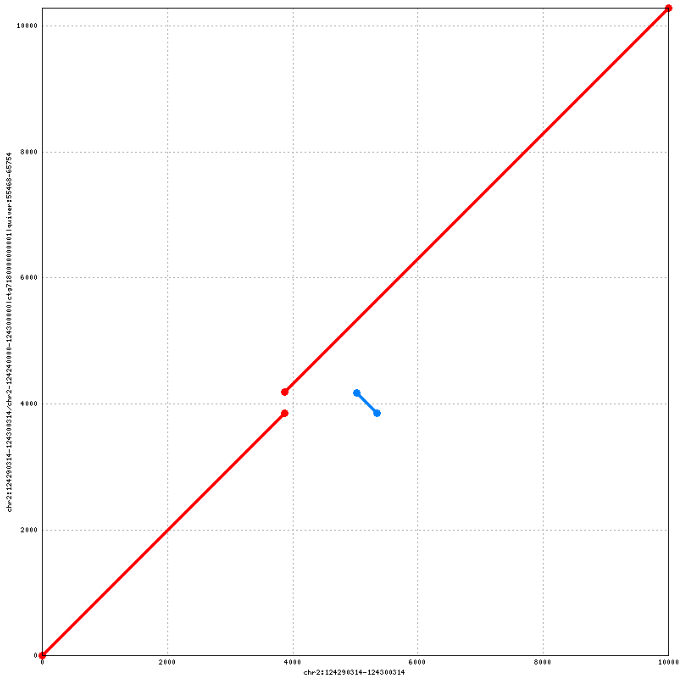


Supplementary Fig. 8: Inverted Duplication predicted within 2:125051481-125053239

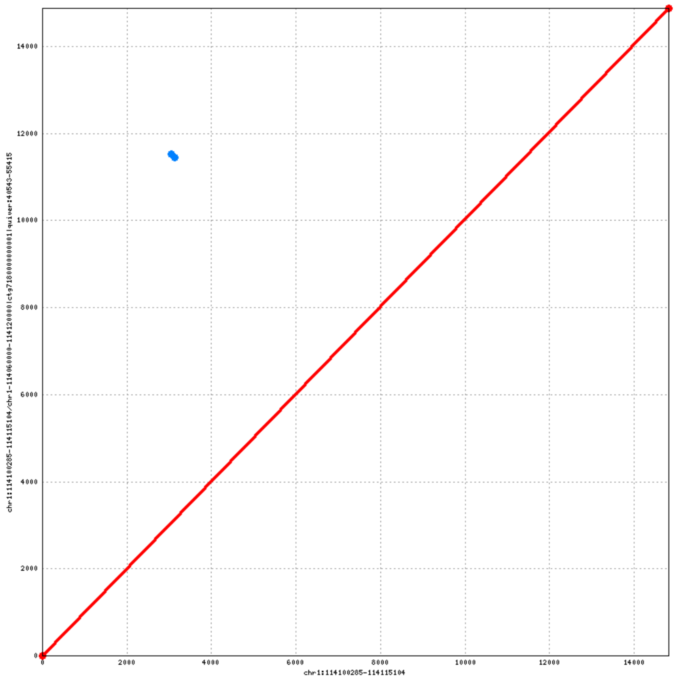


Supplementary Fig. 9: Inverted Duplication predicted within 1:114,645,854-114,654,623.

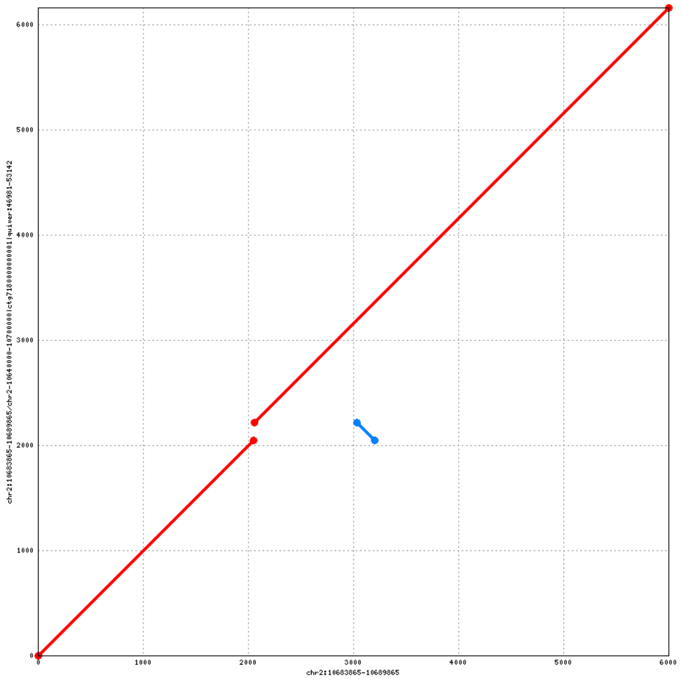


Supplementary Fig. 10: Inverted Duplication predicted within 2:10,825,652-10,827,218.

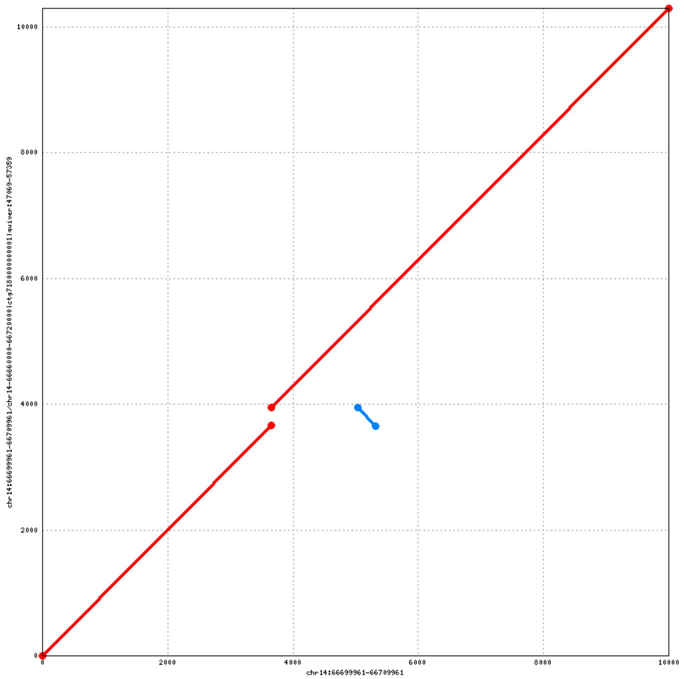


Supplementary Fig. 11: Inverted Duplication predicted within 14:67,169,917-67,171,999.

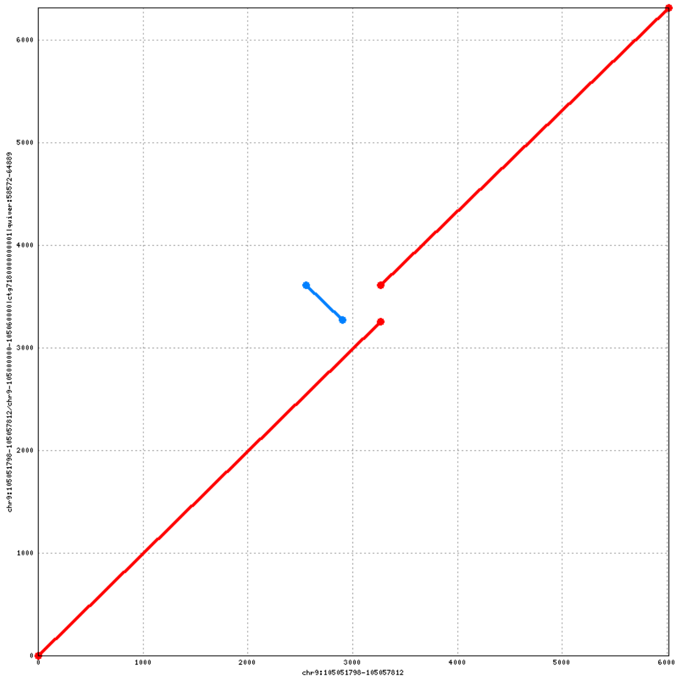


Supplementary Fig. 12: Inverted Duplication predicted within 9:107,816,536-107,817,623.

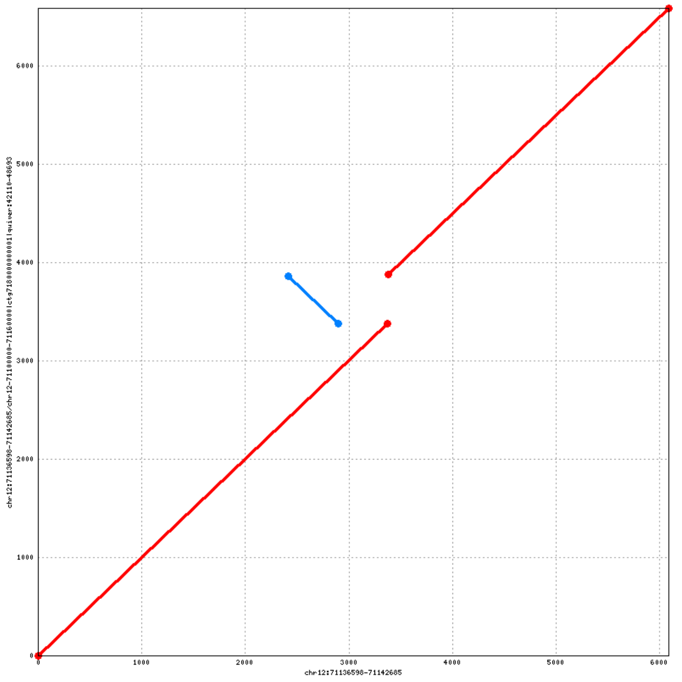


Supplementary Fig. 13: Inverted Duplication predicted within 12:71,532,693-71,534,000.

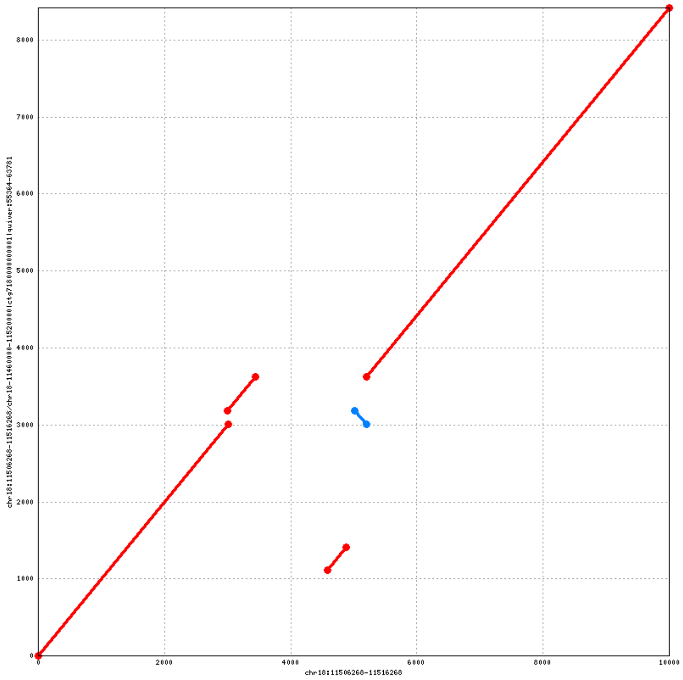


Supplementary Fig. 14: Inverted Duplication predicted within 18:11,508,829-11,511,479.

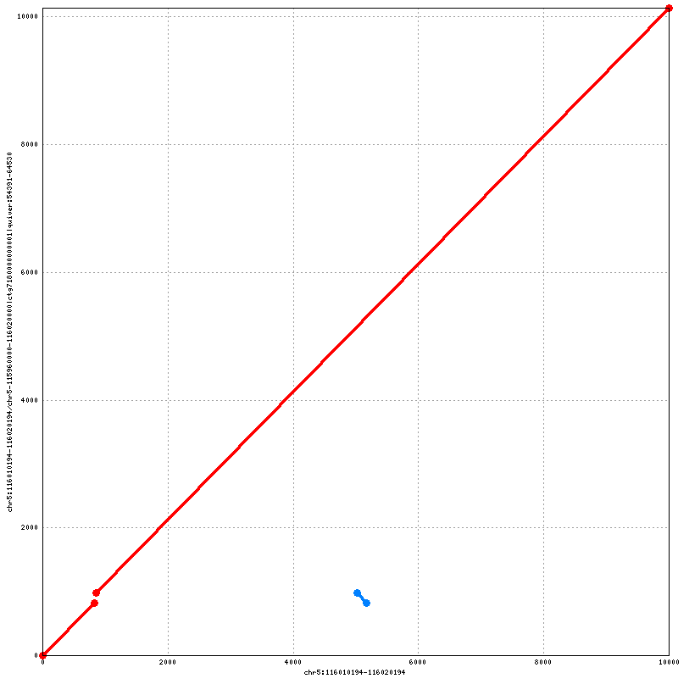


Supplementary Fig. 15: Inverted Duplication predicted within 5:115,346,294-115,351,084.

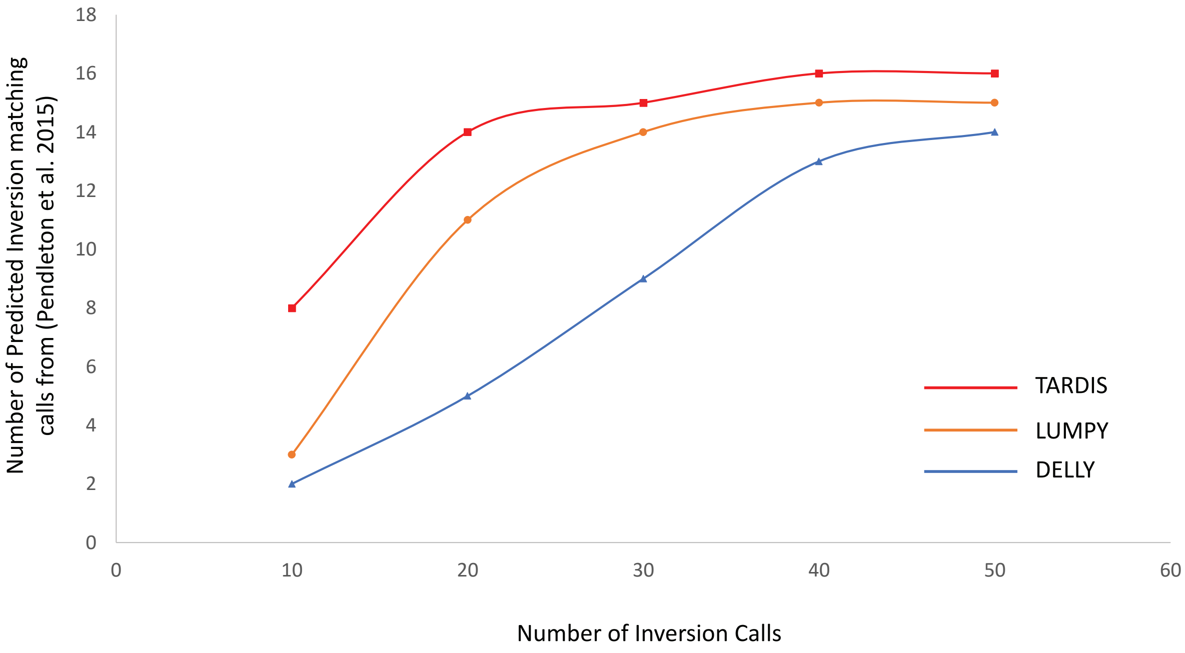


Supplementary Fig. 16: Comparison of top inversion predictions on NA12878 sample against predicted and validated set of inversion of the same samples using PacBio data from [29].

Supplementary Table S4: Performance comparison in terms of time and memory usage for CHM1 and NA12878 genomes.

| | CHM1 | | NA12878 | |
|---|---|---|---|---|
| | Time | Memory | Time | Memory |
| TARDIS | 2h 05m | 9 GB | 3h 56m | 9 GB |
| TIDDIT | 1h 31m | 5 GB | 2h 25m | 5 GB |
| LUMPY | 7h 38m | 7 GB | 11h 05m | 8 GB |
| DELLY | 64h 22m | 7 GB | 33h 05m | 9 GB |
| SoftSV | 175h 26m | 4 GB | 137h 35m | 2 GB |

We benchmarked TARDIS, TIDDIT, LUMPY, DELLY and SoftSV using a haploid (CHM1) and a diploid (NA12878) genome with the same computing resources (Intel(R) Xeon(R) CPU E7- 4830 @ 2.13GHz : 4 CPUs * 8 cores each = 32 cores total 512 GB RAM)
